# Evaluation of morpho-physiological and yield-associated traits of rice (*Oryza sativa* L.) landraces combined with marker-assisted selection under high temperature stress and elevated atmospheric CO_2_ levels

**DOI:** 10.1101/2023.08.31.555684

**Authors:** Merentoshi Mollier, Rajib Roychowdhury, Lanunola Tzudir, Radheshyam Sharma, Bhabesh Gogoi, Prakash Kalita, Devendra Jain, Ranjan Das

**Affiliations:** Department of Crop Physiology, College of Agriculture, Assam Agricultural University, Jorhat 785013, Assam, India; Department of Genetics and Plant Breeding, School of Agricultural Sciences, Nagaland University, Medziphema 797106, Nagaland, India; Department of Plant Pathology and Weed Research, Institute of Plant Protection, Agricultural Research Organization (ARO) - Volcani Institute, Rishon Lezion 7505101, Israel; Department of Agronomy, School of Agricultural Sciences, Nagaland University, Medziphema 797106, Nagaland, India; Biotechnology Centre, Jawaharlal Nehru Krishi Vishwa Vidyalaya, Jabalpur 482004, Madhya Pradesh, India; Department of Soil Sciences, Assam Agricultural University, Jorhat 785013, Assam, India; Department of Molecular Biology & Biotechnology, Rajasthan College of Agriculture, Affiliated to Maharana Pratap University of Agriculture and Technology (MPUAT), Udaipur 313001, Rajasthan, India

**Keywords:** Climate change, Elevated CO_2_, Grain yield, High temperature, Rice landrace, SCoT marker

## Abstract

Rice (*Oryza sativa* L.) has a tremendous domestication history and is presently used as a major cereal all over the world. In Asia, India is considered as one of the centers of origin of *indica* rice and has several native landraces, especially in North-Eastern India (NEI), which have the potential to cope with the negative impact of present-day climate change. The current investigation aimed to evaluate the NEI rice landraces’ potential under high temperatures and elevated CO_2_ levels in comparison with a check variety for phenological, morphological, physiological and yield-associated parameters and molecularly validated with marker-assisted genotyping. The initial experiment was carried out with 75 rice landraces to evaluate their high heat tolerance ability. Seven better-performing landraces along with the check variety (N22) were further evaluated for aforesaid traits across two years (2019 and 2020) under control (or T1) and two stress treatments – (i) mild stress or T2 [CO_2_ 550 ppm + 4° C more than ambient temperature] and (ii) severe stress or T3 [CO_2_ 750 ppm + 6° C more than ambient temperature] using bioreactors. In the molecular analysis, the eight selected genotypes were evaluated through 25 Start Codon Targeted (SCoT) markers. The results revealed that the mild stress (T2) had a positive impact on various morpho-physiological parameters like plant height, number of leaves, leaf area and yield parameters like spikelets panicle^-1^ (S/P), thousand-grain weight (TGW) and grain yield (GY). This effect could be attributed to the genotypes’ ability to maintain a higher photosynthetic rate and possess better tolerance ability to moderately high temperatures. However, under high-temperature conditions in T3, all genotypes exhibited a significant decrease in the studied parameters including GY. It was found that pollen traits were significantly and positively correlated to spikelet fertility% at maturity, which was further significantly associated with GY under applied stress conditions. The physiological traits including shoot biomass were evident to have a significant positive effect on yield-associated parameters like S/P, harvest index (HI), TGW and GY. Overall, two landraces Kohima special and Lisem were found to be better responsive compared to other landraces as well as the check variety N22 under stress conditions. SCoT genotyping amplified a total of 77 alleles out of which 55 were polymorphic with the PIC value ranging from 0.22 to 0.67. The investigation suggests the presence of genetic variation among the tested rice lines and further supports evidence of the closely relatedness of Kohima special and Lisem. These two are two better-performing rice landraces from North-East India based on their improving morpho-physiological parameters and yield attributes in mild and severe high temperature and elevated CO_2_ stress environments. The shortlisted two rice landraces can be used as valuable pre-breeding materials for future rice breeding programs to improve the stress tolerance properties, particularly to high temperatures and elevated CO_2_ levels under ongoing changing climatic scenarios.

## 1. Introduction

Climate change has significant implications for agriculture and food production (Chakraborty et al., 2014). The changes in temperature, precipitation patterns, and extreme weather events associated with climate change can impact various aspects of agricultural systems, including phenology, growth and development, crop yields, and overall food security (Roychowdhury, 2014; Roychowdhury et al., 2020). High temperature and elevated carbon dioxide (CO_2_) levels are two significant consequences of climate change that can have profound impacts on Rice (*Oryza sativa* L.) based agriculture (Hasanuzzaman et al., 2013; Song et al., 2022), which is one of the most important cereals serving as the staple food for approximately 67% of the world population (Senapati et al., 2022). Heatwaves can cause stress to rice plantations by exceeding their temperature tolerance thresholds and lead to reduced photosynthesis, impaired pollen development, and lower crop yields (Hasanuzzaman et al., 2013). Higher CO_2_ concentrations can initially stimulate rice growth due to increased photosynthesis rates which can lead to potentially higher yields (Ainsworth and Long, 2005). However, it is essential to carefully consider the possible advantages of elevated levels of carbon dioxide (CO_2_) in light of the adverse consequences associated with rising temperatures (Kontgis et al., 2019). Based on the current trends, global temperatures can be expected to rise by 3°C at the end of the century, which will lead to fluctuations in climate-related parameters, directly and indirectly impacting on agricultural rice production (Petersen, 2019).

Rice is the world’s most important cereal crop followed by wheat, and provides approximately 50% of calories for almost half of the world’s population and its demand will increase by approximately 28% by 2050 (Zhu et al., 2018). Rice is predominantly cultivated and consumed in Asia – especially in India, China and Japan. India is largely producing *indica* rice (Satapathy et al., 2015). It possesses exceptional genetic diversity, making it one of the few species with such a rich genetic pool on the Indian subcontinent. Through its wide range of domestication history, rice landraces representing traditional and locally adapted varieties, hold immense potential in addressing challenges posed by climate change (Karmakar et al., 2012; Roychowdhury et al., 2013). The North-East (NE) region of India is recognized as a significant hotspot for rice genetic resources due to its diverse rice-growing environments and as a secondary center of origin for rice, the NE Indian region possesses a rich biodiversity of rice germplasm that showcases unique characteristics (Singh and Singh, 2019). As a result of the Green Revolution, most of the ancient rice varieties and landraces were not still cultivated in the majority of the rice growing areas which have been replaced by high-yielding modern cultivars (Roychowdhury et al., 2013). Being the center of rice domestication, the tribal and rural areas of NE India harbour a remarkable rice genetic diversity maintained by farmers and exhibit variations in various traits such as phenology, plant height, photoperiod sensitivity, grain size and shape, aroma, cooking quality, and tolerance to abiotic and biotic stresses (Choudhury et al., 2013). Harnessing the genetic diversity present in these landraces can contribute to the development of new rice varieties that can thrive under changing climate conditions. By incorporating traits like early flowering and maturity, higher pollen vitality and reproductive success, higher biomass, improved plant traits for breeding purposes, and improved grain filling and yield, these landraces offer a promising avenue for sustainable rice production under escalating climate challenges, ultimately bolstering food security for both local communities and the broader population (Kobayasi et al., 2019; Morita et al., 2016; Yamaguchi et al., 2022).

In addition to the breeding values of these landraces under high temperatures and elevated CO_2_ levels, it is also very important to genetic characterization of the landrace population by using molecular genetic markers and to support genotypic performance to screen out better performing landraces for future breeding pipelines. Different molecular markers like start codon targeted (SCoT), simple sequence repeat (SSR), and single nucleotide polymorphic (SNP) markers have been used to characterize rice genotypes and have revealed their profound role in varietal discrimination and trait selection (Roychowdhury et al., 2023a, 2014). SCoT markers are a type of molecular marker used in crop genetic analysis and are based on the polymorphism on short ATG start codon region of genes, making them valuable tools for studying genetic diversity, population structure, and phylogenetic relationships among different rice varieties with stress tolerance properties (Collard and Mackill, 2009). SCoT marker-based analysis presents distinct advantages over other widely used molecular markers like SSR and SNP in rice as it does not require prior knowledge of genomic sequences for primer design, making them particularly valuable for non-model species or less studied rice varieties (Omar et al., 2023). Furthermore, SCoT markers offer higher polymorphism rates compared to SSRs, enabling more accurate discrimination between closely related genotypes, and enhancing the resolution in genetic diversity studies. In comparison to SNP markers, SCoT analysis is cost-effective, as it avoids the resource-intensive processes of SNP discovery and genotyping. SCoT markers also target non-coding regions near start codons, potentially revealing regulatory elements that influence gene expression and functional variations. These features collectively render SCoT marker-based analysis an efficient and versatile tool for studying trait-associated variations in rice, particularly in scenarios where genomic resources are limited or when budget constraints are considered (Rai, 2023).

Hence, the present investigation aims to (a) characterize the native North-East eastern *indica* rice landraces for phenology, morphological, physiological traits, and yield-associated parameters, (b) study the effect of high temperature and elevated CO_2_ on the rice traits for improved yield, and (c) marker-assisted selection of potential rice landraces that are better adapted under aforesaid stress conditions for identification of resilient traits to develop climate-resilient rice varieties.

## 2. Materials and methods

### 2.1 Plant materials

A total of 75 locally grown rice landraces were collected from different rice growing places of Nagaland (Table S1), situated in North-East Indian agro-ecological regions, and were agronomically evaluated in the experimental field (26°45′N latitude, 94°12′E longitude, 87 m altitude above mean sea level) of Assam Agricultural University (AAU), Jorhat, Assam, India during two consecutive rice growing seasons of the year 2019 (Year-1) and 2020 (Year-2). The meteorological data during the cropping seasons (from late June to late December) were obtained from the Department of Agrometeorology, AAU, Jorhat, India and presented graphically in Figure 1.

**Figure 1.**
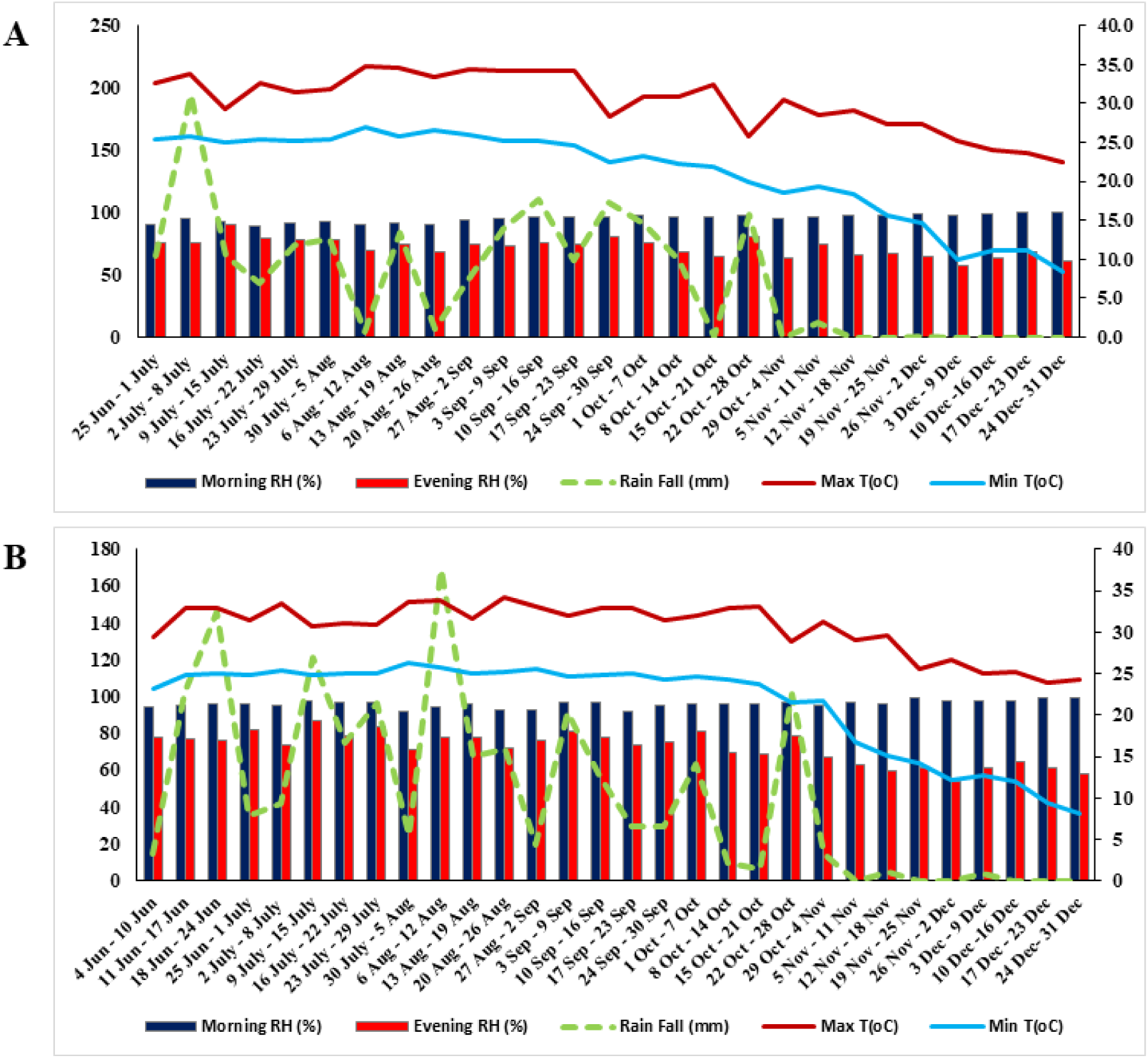
Meteorological data of maximum and minimum temperature (°C), total rainfall (mm), and relative humidity (RH%) in the morning and evening during the cropping seasons (A) Year-1 and (B) Year-2.

### 2.2 Screening the genotypes for high temperature (heat) stress tolerance

All 75 rice landraces were screened for high temperature (heat) tolerance (HT) by the in-field standard Screening Evaluation Score (SES) system of the International Rice Research Institute (IRRI), Philippines (IRRI, 2002). A completely Randomized Design (CRD) with two replications was followed for the field evaluation of seven heat-tolerant genotypes viz. Tatza, Kohima Special, Kakadhan, Laldhan, Tzumma, Lisem and Mapok Temeseng were selected for further detailed evaluation (Table 1). For comparison, one national rice check variety Nagina 22 or N22 was included.

**Table 1.** Shortlisted rice genotypes’ names, collection places from Nagaland state of India and genotypic types.

| Accession No. | Rice genotypes | Places of collection | District | State | Genotypic type |
| --- | --- | --- | --- | --- | --- |
| G1 | Tatza | Punglao | Dimapur | Nagaland | Landrace |
| G2 | Kohima Special | Molva | Dimapur | Nagaland | Landrace |
| G3 | Kaladhan | Jharnapani | Dimapur | Nagaland | Landrace |
| G4 | Laldhan | Medziphema | Dimapur | Nagaland | Landrace |
| G5 | Tzumma | Sochunoma | Dimapur | Nagaland | Landrace |
| G6 | Lisem | Mopongchuket | Mokokchung | Nagaland | Landrace |
| G7 | Mapok Temeseng | Merangkong | Mokokchung | Nagaland | Landrace |
| G8 | Nagina 22<br>(or N22) | National Rice Research Institute<br>(NRRI),<br>Cuttack, India | Bidyadharpur | Orissa | Improved check variety |

### 2.3 Experimental set-up and treatments of high temperature and elevated CO_2_ stress

Surface sterilized (with 0.1% sodium hypochlorite) rice seeds of selected eight genotypes were sown in plastic cups containing rice field soil with organic matter (50:50) and grown for 30 days. Then 10 healthy plantlets were transferred to a bigger pot (30 cm^3^) in five replications with the same soil mixture for each treatment and maintained in the bioreactors (Genesis Technologies, Maharashtra, India) established in the Stress Physiology Laboratory, Department of Crop Physiology (AAU, Jorhat, India). Such bioreactors were used for further study with high-temperature and elevated CO_2_ stress treatments. Factorial Completely Randomized Design (FCRD) with two factors (two bioreactors per treatment) and 5 replicates have been employed for the stress treatments. Treatment-1 (T1) is the control set and maintained the plant pots under ambient temperature and CO_2_, whereas Treatment-2 (T2) comprises CO_2_ (550±20ppm) with + temperature (ambient + 4°C) and Treatment-3 (T3) comprises CO_2_ (750±20ppm) with + temperature (ambient + 6°C), maintained in individual bioreactors as follows:

T1: Control (ambient CO_2_ and temperature)

T2 (mild-stress): CO_2_ (550±20ppm) with + temperature (ambient + 4°C)

T3 (severe-stress): CO_2_ (750±20ppm) with + temperature (ambient + 6°C)

### 2.4 Growth conditions in bioreactors

Inside the bioreactor, the CO_2_ level was maintained throughout the rice plant growth during 09:00 – 14:00 daily. The elevation of temperature was maintained from the tillering stage to just before maturity through an Infra-Red (IR) heater regulated by SCADA software. Temperature transmitters based on Resistance Temperature Detectors (RTDs) sensors were placed in each chamber to get data in the control room through four core-shielded cables. IR heaters are fitted inside the bioreactors to elevate the temperature and air conditioners are placed inside which are operated with the help of a remote controller. All the data were recorded in the computer placed in the control room.

### 2.5 Phenotyping of phenological, morpho-physiological and yield-associated traits

All the agronomic practices were followed during the growth condition as per the desired standard. Phenological parameters like days to flowering (DtoF) and days to physiological maturity (DtoPM) were recorded at particular growth stages. The former represents the number of days taken from sowing to the initiation of 50% flowering of panicles in a pot. The physiological maturity is expressed as the number of days taken from sowing to 70-80% crop maturity (late grain filling stage). Canopy temperature or CT (°C) is measured by the infrared thermometer (IRT) within six hours after treatment. Agro-morphological parameters such as number of tillers plant^-1^ (TL/P), leaf area or LA (cm^2^), leaf area index (LAI), relative leaf water content (RLWC), root length or RL (cm) and volume or RV (cm^3^), root biomass or RB (g), anther length (AnL) and pollens anther^-1^ (P/A) were recorded during the grain filling stage. The number of tillers per hill (main plant) was counted on the tagged plants in the pot. The numbers of green leaves (LN) and LA were measured using a leaf area meter (LI 3000). LAI expresses the ratio of leaf surface to the ground area occupied by the plant as per (Evans, 1972). RLWC was calculated as per the formula used by (Barrs and Weatherley, 1962) using twenty-leaf discs. Soil-Plant Analysis Development (SPAD) chlorophyll meter (SPAD-502, Osaka, Japan) was used to obtain SPAD values (SPAD units) from the four uppermost fully expanded leaves on each plant during grain filling stage as per (Frankin et al., 2020). Normalized difference vegetation index (NDVI) was recorded at the same time using GreenSeeker handheld sensor (NTech Industry Inc., Ukiah, CA) as per (Kimaro et al., 2023). For root characteristics, a set of plants was uprooted very carefully during maturity and roots were separated from above-ground parts, and evaluated RL. RV was determined by the water displacement method used by (Raja and Bishnoi, 1990). Then the roots were dried in an oven at 80°C for 3 days and the RB was recorded using an electronic weight machine (Shimadzu ELB600, Japan). AnL (as mm) and P/A were calculated at the time of flowering after staining the anthers with safranine and observed under a light microscope (Das and Das, 2021). Plant height or PH (as cm) was measured at maturity with the help of a meter scale from the base of the plant to the apex of the main axis. Yield components such as the number of spikes panicle^-1^ (SP/P), spikelet fertility^-1^ (SF%), number of filled grains panicle^-1^ (G/P), thousand-grain weight (TGW as g), grain yield (GY as g plant^-1^) and harvest index (HI) were measured at maturity. During maturity, all above-ground plant parts (vegetative + reproductive) of each plant with tillers per pot were manually harvested as shoot biomass (SB) and separated vegetative parts from the panicle. Five random panicles were collected from each pot and the number of spikes and filled grains were counted for each panicle. The spikelet fertility was estimated as the ratio of the number of filled grains to the total number of florets. Then the grains were dried and threshed manually. The number of grains and their weight (GY and TGW) were measured using an electronic weight machine (Shimadzu ELB600, Japan). HI was measured as the proportion of GY to the total above-ground dry matter per pot and expressed in per cent (%).

### 2.6 Genotyping using Start Codon Targeted (SCoT) markers

Genomic DNA was isolated from the young and healthy rice leaves during the maturity stage following a pre-standardized protocol (Roychowdhury et al., 2012). The DNA quality was checked using 0.8% agarose gel electrophoresis and nanodrop reading (Ganie et al., 2014). The high-quality DNA was used for polymerase chain reaction (PCR) based genotyping using a total of 30 Start Codon Targeted (SCoT) markers and carried out the PCR reaction in a thermal cycler (M. J. Research, MC 013130) as performed by (Agarwal et al., 2008) to check the polymorphic ability and discard the monomorphic SCoT markers from the analysis. The list of 25 polymorphic SCoT primers used in the present investigation is presented in Table S2. The PCR amplified products were resolved in 1.5% agarose gel following the protocol given by (Ganie et al., 2016), stained by ethidium bromide (EtBr) and documented in a UV gel documentation system (Perkin Elmer, Geliance 200 imaging system). The length of the amplified DNA bands was determined with reference to the 100 bp DNA ladder (Fermentus Life Sci. USA) included in the gel as a size marker. The molecular weight (nucleotide base pairs) of the most intensely amplified bands for each microsatellite marker was analyzed using the software AlphaEaseFC (v4.0, USA) as reported by (Roychowdhury et al., 2013). The electrophoretic banding pattern generated from SCoT primers was used to calculate pair-wise genetic similarity of rice genotypes and a dendrogram was constructed by using unweighted pair group analysis (UPGMA) using NTSYS-pc version 2.02e software.

### 2.7 Statistical analyses

Statistical analysis was done using JMP v16.0 (SAS, NC, USA) software and R-programming for Windows (x64). Differences between the treatments were analyzed using Tukey-Kramer post-hoc test. Analysis of variance (ANOVA) and genotype-treatment-year interaction (GxTxY) has been done using a mixed model with blocks having random effects. Correlations between the parameters of the control set have been evaluated using the corrplot package of R v4.0.3. Principal component analysis (PCA) was graphically computed using the first two PC1 and PC2 through multivariate analysis.

## 3. Results

A total of 75 rice landraces collected from different places of Nagaland (NEI region) were initially screened for high-temperature (heat) stress tolerance based on gradual heat responsive phenotypic changes of leaf i.e. days to stay healthy, leaf folding and rolling, leaf tip and margin drying, whole leaf drying and plant death (Table S1). After such initial screening, seven landraces [Tatza (G1), Kohima special (G2), Kaladhan (G3), Laldhan (G4), Tzumma (G5), Lisem (G6) and Mapok Temeseng (G7)] were found high temperature tolerant and were selected with comparing to the national check variety – N22 for further study (Table 1) to describe the interactive effects of control (T1 – no stress imposed) and two different stress level – mild (T2) and severe (T3) with high temperatures and elevated CO_2_. In both control and two stress conditions, phenological (DtoF, DtoPM), morphological (PH, LN, RL, RV, RB, SB, AnL, P/A), physiological (CT, SPAD, NDVI, RLWC, LA) and yield-associated traits (GY, HI, TGW, G/P, SP/P, panicle length or PL, SF%) were evaluated in two years (2019 and 2020).

### 3.1 Effect of high temperatures and elevated CO_2_ level on rice phenology, morpho-physiological and yield-associated parameters

The diversity in the phenological, morphological, physiological and yield-associated traits have been detected in control and stress treatments across the two years (2019 and 2020) of study on the selected eight rice genotypes and the mean values are shown in Table 2. DtoF, RB m^-2^, TL m^-2^ and PL have been found to have the least effect in different treatments. When comparing similar treatments in both studied years for each trait, variable performance is almost stable without much deviation. A clear trend was observed which indicating that mild stress treatment (T2) yielded better variable values than the control across two years, which drastically declined in the severe stress (T3) of high temperature and elevated CO_2_ levels. Reduction in DtoF and DtoPM in T2 for the studied genotypes is vital for the shortening of life span which may correspond to high yielding under stress conditions. SPAD (chlorophyll content) values were decreased thoroughly from control to T2 and then T3 in both years. But the NDVI shows such a similar trend only for year-1. In year-2, NDVI raised little in T2 compared to the control, but those were non-significant (p<0.05 in Tukey-Kramer post-hoc test). Almost it has been observed that the trait values in T3 were generally much lesser than the values in control, but for some traits it was higher than the control, e.g. CT, root traits (RL, RV, RB) and PL.

**Table 2.**
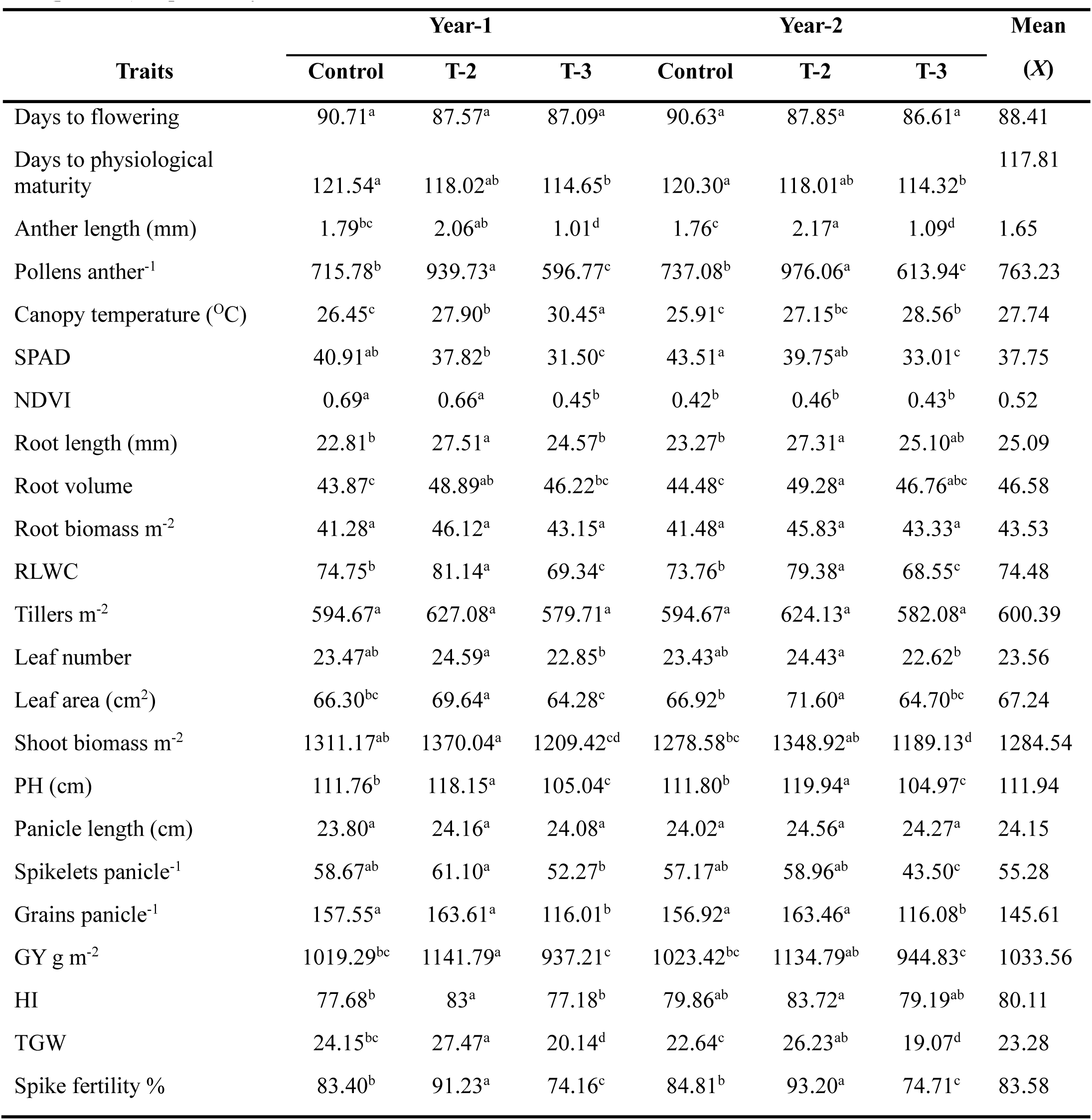
Mean values of the studied traits under control and two different stress treatments in two consecutive years – Year-1 and Year-2. The mean values with different letters are significantly different (p = 0.05) as per Tukey-Kramer’s LSD test.

| Traits | Year-1 |  |  | Year-2 |  |  | Mean<br>(X) |
| --- | --- | --- | --- | --- | --- | --- | --- |
|  | Control | T-2 | T-3 | Control | T-2 | T-3 |  |
| Days to flowering | 90.71<sup>a</sup> | 87.57<sup>a</sup> | 87.09<sup>a</sup> | 90.63<sup>a</sup> | 87.85<sup>a</sup> | 86.61<sup>a</sup> | 88.41 |
| Days to physiological maturity | 121.54<sup>a</sup> | 118.02<sup>ab</sup> | 114.65<sup>b</sup> | 120.30<sup>a</sup> | 118.01<sup>ab</sup> | 114.32<sup>b</sup> | 117.81 |
| Anther length (mm) | 1.79<sup>bc</sup> | 2.06<sup>ab</sup> | 1.01<sup>d</sup> | 1.76<sup>c</sup> | 2.17<sup>a</sup> | 1.09<sup>d</sup> | 1.65 |
| Pollens anther<sup>-1</sup> | 715.78<sup>b</sup> | 939.73<sup>a</sup> | 596.77<sup>c</sup> | 737.08<sup>b</sup> | 976.06<sup>a</sup> | 613.94<sup>c</sup> | 763.23 |
| Canopy temperature (°C) | 26.45<sup>c</sup> | 27.90<sup>b</sup> | 30.45<sup>a</sup> | 25.91<sup>c</sup> | 27.15<sup>bc</sup> | 28.56<sup>b</sup> | 27.74 |
| SPAD | 40.91<sup>ab</sup> | 37.82<sup>b</sup> | 31.50<sup>c</sup> | 43.51<sup>a</sup> | 39.75<sup>ab</sup> | 33.01<sup>c</sup> | 37.75 |
| NDVI | 0.69<sup>a</sup> | 0.66<sup>a</sup> | 0.45<sup>b</sup> | 0.42<sup>b</sup> | 0.46<sup>b</sup> | 0.43<sup>b</sup> | 0.52 |
| Root length (mm) | 22.81<sup>b</sup> | 27.51<sup>a</sup> | 24.57<sup>b</sup> | 23.27<sup>b</sup> | 27.31<sup>a</sup> | 25.10<sup>ab</sup> | 25.09 |
| Root volume | 43.87<sup>c</sup> | 48.89<sup>ab</sup> | 46.22<sup>bc</sup> | 44.48<sup>c</sup> | 49.28<sup>a</sup> | 46.76<sup>abc</sup> | 46.58 |
| Root biomass m<sup>-2</sup> | 41.28<sup>a</sup> | 46.12<sup>a</sup> | 43.15<sup>a</sup> | 41.48<sup>a</sup> | 45.83<sup>a</sup> | 43.33<sup>a</sup> | 43.53 |
| RLWC | 74.75<sup>b</sup> | 81.14<sup>a</sup> | 69.34<sup>c</sup> | 73.76<sup>b</sup> | 79.38<sup>a</sup> | 68.55<sup>c</sup> | 74.48 |
| Tillers m<sup>-2</sup> | 594.67<sup>a</sup> | 627.08<sup>a</sup> | 579.71<sup>a</sup> | 594.67<sup>a</sup> | 624.13<sup>a</sup> | 582.08<sup>a</sup> | 600.39 |
| Leaf number | 23.47<sup>ab</sup> | 24.59<sup>a</sup> | 22.85<sup>b</sup> | 23.43<sup>ab</sup> | 24.43<sup>a</sup> | 22.62<sup>b</sup> | 23.56 |
| Leaf area (cm<sup>2</sup>) | 66.30<sup>bc</sup> | 69.64<sup>a</sup> | 64.28<sup>c</sup> | 66.92<sup>b</sup> | 71.60<sup>a</sup> | 64.70<sup>bc</sup> | 67.24 |
| Shoot biomass m<sup>-2</sup> | 1311.17<sup>ab</sup> | 1370.04<sup>a</sup> | 1209.42<sup>cd</sup> | 1278.58<sup>bc</sup> | 1348.92<sup>ab</sup> | 1189.13<sup>d</sup> | 1284.54 |
| PH (cm) | 111.76<sup>b</sup> | 118.15<sup>a</sup> | 105.04<sup>c</sup> | 111.80<sup>b</sup> | 119.94<sup>a</sup> | 104.97<sup>c</sup> | 111.94 |
| Panicle length (cm) | 23.80<sup>a</sup> | 24.16<sup>a</sup> | 24.08<sup>a</sup> | 24.02<sup>a</sup> | 24.56<sup>a</sup> | 24.27<sup>a</sup> | 24.15 |
| Spikelets panicle<sup>-1</sup> | 58.67<sup>ab</sup> | 61.10<sup>a</sup> | 52.27<sup>b</sup> | 57.17<sup>ab</sup> | 58.96<sup>ab</sup> | 43.50<sup>c</sup> | 55.28 |
| Grains panicle<sup>-1</sup> | 157.55<sup>a</sup> | 163.61<sup>a</sup> | 116.01<sup>b</sup> | 156.92<sup>a</sup> | 163.46<sup>a</sup> | 116.08<sup>b</sup> | 145.61 |
| GY g m<sup>-2</sup> | 1019.29<sup>bc</sup> | 1141.79<sup>a</sup> | 937.21<sup>c</sup> | 1023.42<sup>bc</sup> | 1134.79<sup>ab</sup> | 944.83<sup>c</sup> | 1033.56 |
| HI | 77.68<sup>b</sup> | 83<sup>a</sup> | 77.18<sup>b</sup> | 79.86<sup>ab</sup> | 83.72<sup>a</sup> | 79.19<sup>ab</sup> | 80.11 |
| TGW | 24.15<sup>bc</sup> | 27.47<sup>a</sup> | 20.14<sup>d</sup> | 22.64<sup>c</sup> | 26.23<sup>ab</sup> | 19.07<sup>d</sup> | 23.28 |
| Spike fertility % | 83.40<sup>b</sup> | 91.23<sup>a</sup> | 74.16<sup>c</sup> | 84.81<sup>b</sup> | 93.20<sup>a</sup> | 74.71<sup>c</sup> | 83.58 |

It is shown that anther length and pollens anther^-1^ are significantly contribute to higher SF% at maturity in control and treatments. T2 is much more productive than the control, but severe stress (T3) is reducing dramatically SF% (Figure 2A-B). In the associationship between the AL and SF%, the genotypic variation in T3 is much more distinct rather than the control and T2, which is quite overlapping with P/A. Also, SF% positively and significantly influences GY (Figure 2C). There is a clear association of SB with major yield attributes i.e. PL, SP/P, G/P, GY, HI and TGW has been found (Figure 2D-I). SB is mostly positively and significantly associated with all the yield attributes, except PL where the association is non-significant with a very low regression (R^2^) value (Figure 2D). SB is highly influencing GY with higher significance levels with higher R^2^ values across the treatments (Figure 2G).

**Figure 2.**
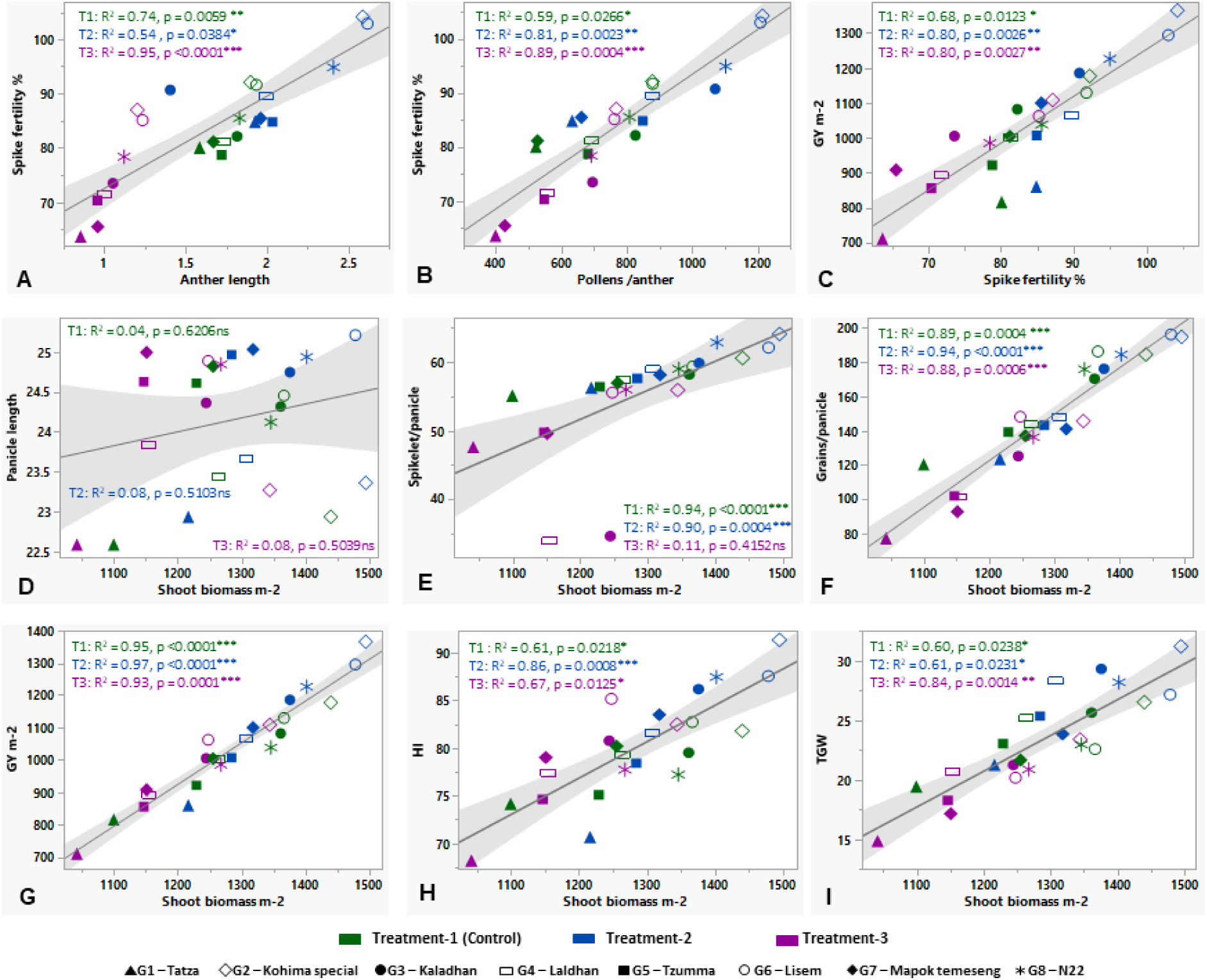
Relationship of pollen traits, biomass and yield-associated parameters in studied rice genotypes under control and stress treatments (mild and severe). ‘*’, ‘**’ and ‘***’ denote the significance level at p ≤ 0.05, p ≤ 0.01 and p ≤ 0.001, respectively. ‘ns’ = non-significant.

It has been found that stress-responsive physiological traits like CT, SPAD, NDVI, RL, RLWC and LA are significantly associated with GY m^-2^ across the control (T1) and two stress treatments – mild (T2) and severe (T3) (Figure 3A-F). CT is negatively related to the GY increment, but the rest of the physiological traits show a positive association with GY. Control and the stress treatments are significant in all trait-association cases, except the control between RLWC and GY with a comparatively low R^2^ value i.e. 0.42 (Figure 3E).

**Figure 3.**
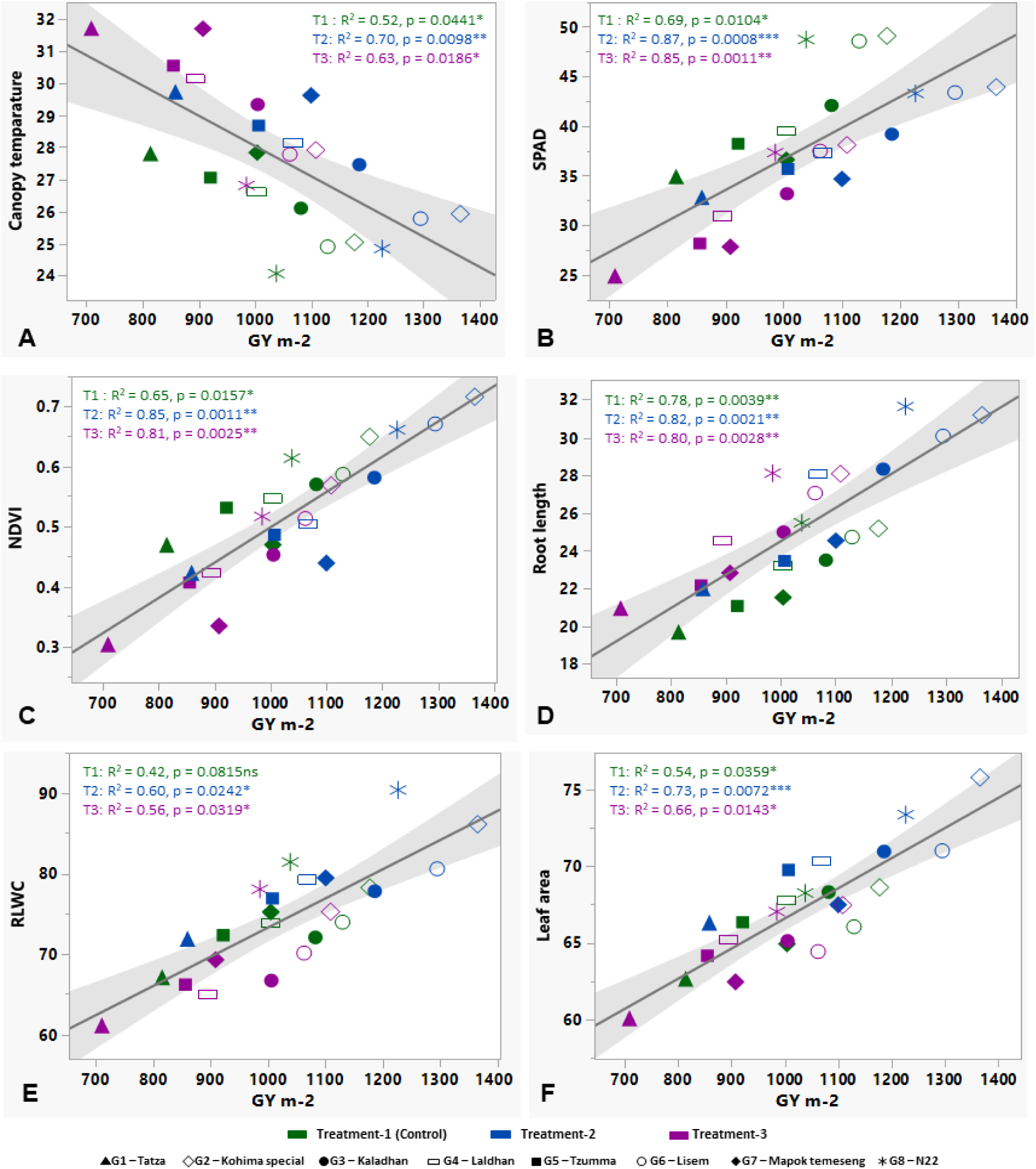
Relationship of grain yield (GY m^-2^) with the stress-responsive physiological parameters – canopy temperature (A), SPAD (B), NDVI (C), root length (D), relative leaf water content or RLWC € and leaf area in cm^2^ (F) under control and two different stress conditions (mild and severe). ‘*’, ‘**’ and ‘***’ denote the significance level at p ≤ 0.05, p ≤ 0.01 and p ≤ 0.001, respectively. ‘ns’ = non-significant.

### 3.2 Inter-relationship between the studied traits

Variability of the studied traits has been revealed through their inter-relationship by considering the control (T1) and the corrplot is shown in Figure 4A-B. In both years (year-1 and year-2), DtoPM, PL, SP/P and TGW are greatly found non-significant with other traits. CT is shown mostly negatively correlated with other traits. The most positive correlation is provided by AnL, P/A, SPAD, NDVI, RL, LN, SB, SP/P, G/P, GY and SF%. In year 1, the highest positive significant correlation (r=0.94, p<0.001) is shown between NDVI and G/P and between P/A and GY (Figure 4A); whereas in year-2, the same highest correlation was observed for P/A associated with LA and TGW (Figure 4B).

**Figure 4.**
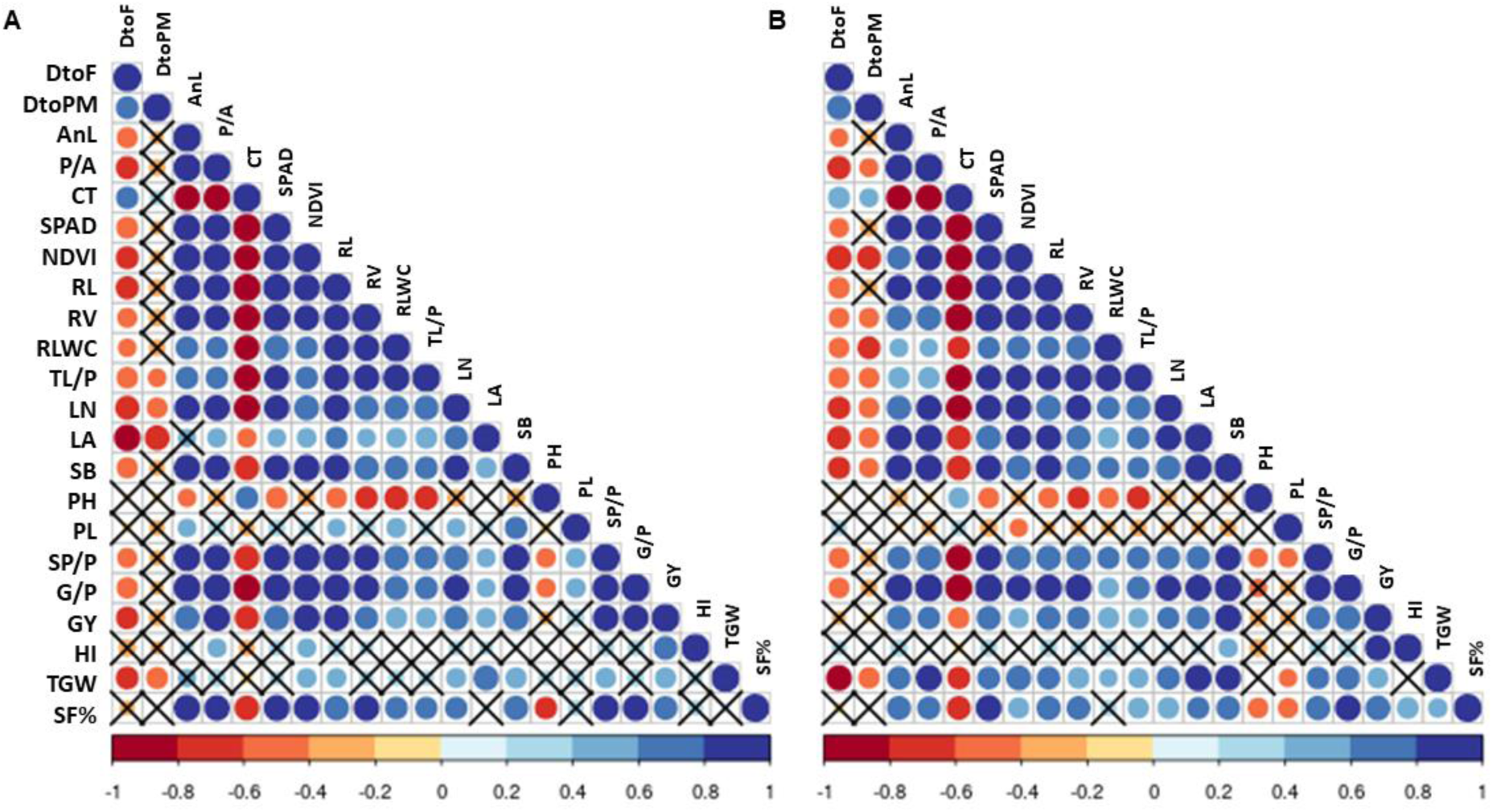
Correlation matrix in the form of corrplot using the studied parameters of rice genotypes of control set in both year 1 (A) and year 2 (B). The range of correlation coefficient ® is shown in the horizontal index bar. The cross mark (X) denotes the non-significant correlation (p=0.05). DtoF – Days to flowering, DtoPM – Days to physiological maturity, AnL – anther length, P/A – pollens per anther, CT – canopy temperature, SPAD – Soil-Plant Analysis Development, NDVI – Normalized difference vegetation index, RL – root length, RV – root volume, RLWC – relative leaf water content, TL/P – tillers per plant, LN – leaf number, LA – leaf area, SB – shoot biomass, PH – plant height, PL – panicle length, SP/P – spikelets per panicle, G/P – grains per panicle, GY – grain yield, HI – harvest index, TGW – thousand-grain weight, SF% - spikelet fertility %.

### 3.3 Combinatorial interaction between genotypes (G), treatments (T) and years (Y)

Analysis of variance (ANOVA) of the studied parameters of the selected eight rice genotypes in two consecutive years shows enormous variability for genotypes (G), treatments (T), years (Y) and their all combinatorial interactions (GxT, GxY, TxY, GxTxY) and shown in Table 3. ANOVA shows that R^2^ of the traits ranged between 0.76 (PL) and 0.99 (DtoF, P/A, CT, NDVI, RB, RLWC, G/P and SF%). Most of all traits show significant differences in the case of genotypes, treatments and years individually, but in some instances, years do not produce significant variable differences, e.g. phenology (DtoF, DtoPM), AnL, RB, TL m^-2^ and LN. A significant difference for GxTxY was observed in a few traits like AnL, P/A, CT, NDVI, SP/P, G/P and SF%. In all combinatorial aspects, CT, NDVI, SF%, P/A, SPAD, LA, SP/P and G/P were producing the maximum significant variation of the factor combinations.

**Table 3.** Analysis of variance (ANOVA) of the studied parameters of rice genotypes with genotypes (G), treatments (T) and years (Y) interaction.

| Fixed factors | DF | Studied parameters |  |  |  |  |  |
| --- | --- | --- | --- | --- | --- | --- | --- |
|  |  | DtoF | DtoPM | AnL | P/A | CT | SPAD |
| G | 7 | <0.0001*** | <0.0001*** | <0.0001*** | <0.0001*** | <0.0001*** | <0.0001*** |
| T | 2 | <0.0001*** | <0.0001*** | <0.0001*** | <0.0001*** | <0.0001*** | <0.0001*** |
| Y | 1 | 0.7054 | 0.035* | 0.1941 | <0.0001*** | <0.0001*** | <0.0001*** |
| GxT | 14 | 0.1518 | 0.0888 | <0.0001*** | <0.0001*** | <0.0001*** | <0.0001*** |
| GxY | 7 | 0.0544 | <0.0001*** | 0.0139* | <0.0001*** | <0.0001*** | <0.0001*** |
| TxY | 2 | 0.4244 | 0.1119 | 0.3264 | 0.1361 | <0.0001*** | 0.0312* |
| GxTxY | 14 | 0.9984 | 0.9711 | 0.0057** | <0.0001*** | <0.0001*** | 0.9573 |
| $R^2$ | | 0.99 | 0.96 | 0.89 | 0.99 | 0.99 | 0.98 |
| Fixed factors | DF | NDVI | RL | RV | RB | RLWC | TL/P |
| G | 7 | <0.0001*** | <0.0001*** | <0.0001*** | <0.0001*** | <0.0001*** | <0.0001*** |
| T | 2 | <0.0001*** | <0.0001*** | <0.0001*** | <0.0001*** | <0.0001*** | <0.0001*** |
| Y | 1 | <0.0001*** | <0.0001*** | <0.0001*** | 0.732 | <0.0001*** | 0.9692 |
| GxT | 14 | <0.0001*** | <0.0001*** | <0.0001*** | <0.0001*** | <0.0001*** | 0.0372* |
| GxY | 7 | <0.0001*** | 0.0057** | 0.0313* | 0.8495 | <0.0001*** | 0.0394* |
| TxY | 2 | <0.0001*** | 0.0418* | 0.7323 | 0.0812 | 0.0038** | 0.8749 |
| GxTxY | 14 | <0.0001*** | 0.99 | 0.996 | 0.994 | 0.4069 | 0.9938 |
| $R^2$ | | 0.99 | 0.97 | 0.98 | 0.99 | 0.99 | 0.97 |
| Fixed factors | DF | LN | LA | LAI | SB | PH | PL |
| G | 7 | <0.0001 | <0.0001*** | <0.0001*** | <0.0001*** | <0.0001*** | <0.0001*** |
| T | 2 | <0.0001 | <0.0001*** | <0.0001*** | <0.0001*** | <0.0001*** | 0.0087* |
| Y | 1 | 0.1831 | <0.0001*** | <0.0001*** | <0.0001*** | 0.0067** | 0.0218* |
| GxT | 14 | 0.5779 | <0.0001*** | <0.0001*** | <0.0001*** | <0.0001*** | 0.9957 |
| GxY | 7 | 0.0099** | <0.0001*** | <0.0001*** | 0.9057 | 0.1162 | <0.0001*** |
| TxY | 2 | 0.7737 | <0.0001*** | 0.6975 | 0.2438 | 0.0006*** | 0.6283 |
| GxTxY | 14 | 0.9998 | 0.7161 | 0.7323 | 0.9393 | 0.9945 | 0.9403 |
| $R^2$ | | 0.92 | 0.96 | 0.98 | 0.98 | 0.97 | 0.76 |
| Fixed factors | DF | SP/P | G/P | GY | HI | TGW | SF% |
| G | 7 | <0.0001 | <0.0001*** | <0.0001*** | <0.0001*** | <0.0001*** | <0.0001*** |
| T | 2 | <0.0001 | <0.0001*** | <0.0001*** | <0.0001*** | <0.0001*** | <0.0001*** |
| Y | 1 | <0.0001 | <0.0001*** | <0.0001*** | <0.0001*** | <0.0001*** | <0.0001*** |
| GxT | 14 | <0.0001 | <0.0001*** | <0.0001*** | <0.0001*** | <0.0001*** | <0.0001*** |
| GxY | 7 | 0.0012** | <0.0001*** | <0.0001*** | <0.0001*** | <0.0001*** | <0.0001*** |
| TxY | 2 | 0.0064** | 0.2735 | 0.3247 | 0.2719 | 0.1851 | 0.0035** |
| GxTxY | 14 | <0.0001*** | <0.0001*** | 0.8294 | 0.6204 | 0.2886 | 0.0418* |
| $R^2$ | | 0.77 | 0.99 | 0.98 | 0.90 | 0.98 | 0.99 |

G – genotype, T – treatments, Y – years, DtoF – Days to flowering, DtoPM – Days to physiological maturity, AnL – anther length, P/A – pollens per anther, CT- canopy temperature, SPAD, NDVI, RL – root length, RV – root volume, RB – root biomass, RLWC – relative leaf water content, TL/P – tillers per plant, LN – leaf number, LA – leaf area, LAI – leaf area index, SB – shoot biomass, PH – plant height, PL – panicle length, SP/P – spikelets per panicle, G/P – grains per panicle, GY – grain yield, HI – harvest index, TGW – thousand-grain weight, SF% - spikelet fertility %. ‘\*’, ‘\*\*’ and ‘\*\*\*’ denote the significance level at $p \leq 0.05$ , $p \leq 0.01$ and $p \leq 0.001$ , respectively.

### 3.4 Multivariate analysis of the phenotypic parameters

Principal component analysis in two consecutive years exhibited a clear separation of treatments with distinct trait vectors. In year-1 (Figure 5A1, A2), the rice genotypes were plotted with PC1 explaining 62.4% of the phenotypic variation. PC1 is positively loaded with plant traits (NDVI, SB, RLWC, LN, RL, TL/P) and yield attributes (SF%, G/P, TGW, GY, HI) and negatively loaded with PH and PL. Similarly, PC2 contributes 14.8% and is positively loaded with phenology (DtoF and DtoPM), and negatively loaded with CT. In the second year (Figure 5B1, B2), almost the vector trend and genotypic position are similar and static. PC1 and PC2 are explaining 59.9% and 18.4% of trait variation, respectively. PC1 is positively loaded with plant traits (NDVI, RLWC, LN, TL/P, RL) and yield attributes (SF%, G/P, GY, TGW) and negatively loaded to PH and SP/P. On the other hand, PC2 is contributing in the same way in year 1. Genotypic discrimination as per the treatment is very clear in the PCA chart which shows treatment-3 (severe high temperature and CO_2_ stress) is much more distinctive from the control and Treatment-2 (severe high temperature and CO_2_ stress). Although, control and treatment-2 are almost mixed and not distinctive between the genotypes.

**Figure 5.**
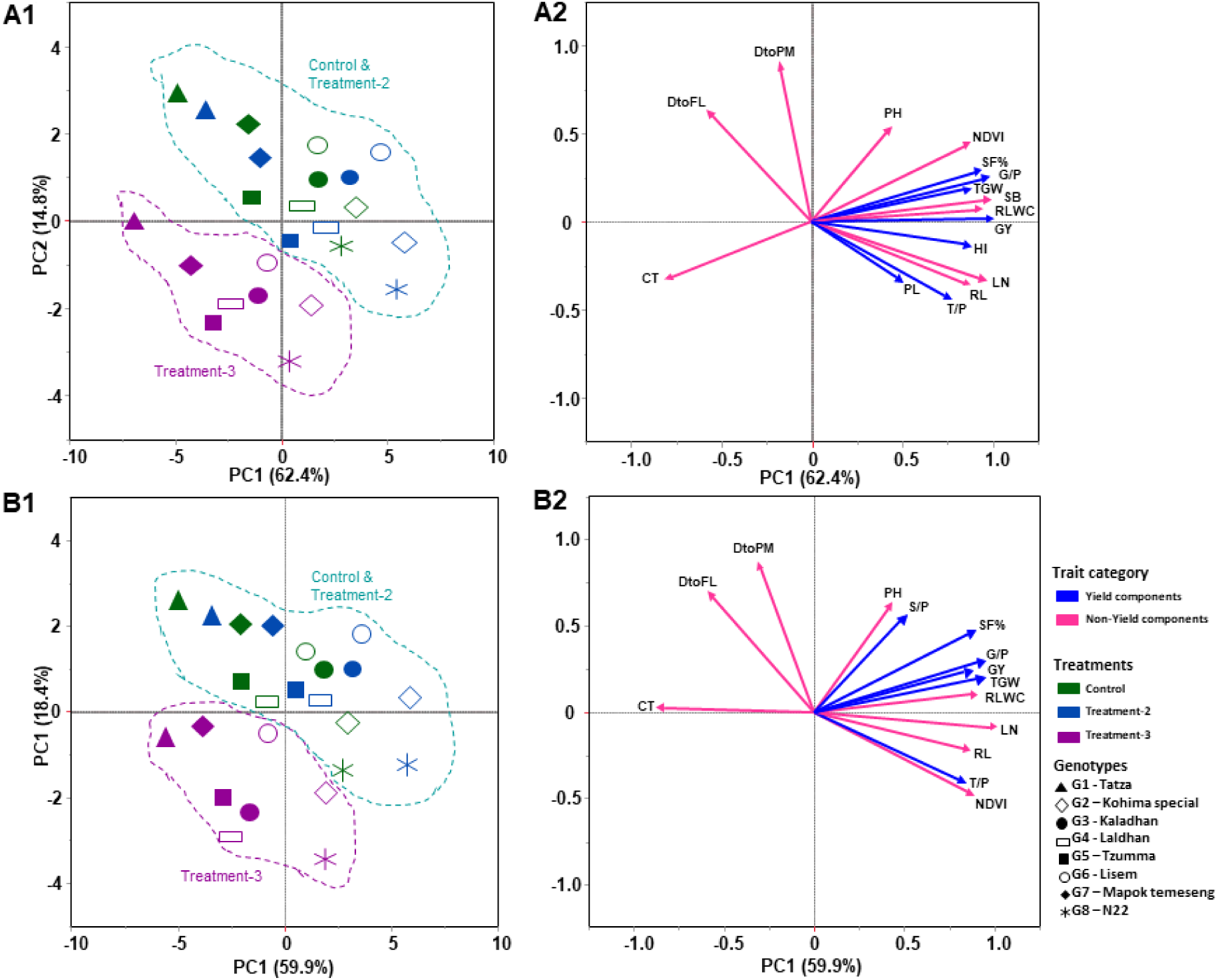
Principal component analysis (PCA) for the studied parameters of yield and non-yield components of eight rice genotypes for consecutive two years of growing seasons under control and two treatment (treatment-2 and treatment-3) conditions. Control (ambient temparature and CO2), treatment 2 (ambient temperature + 4°C, CO2 550±20ppm) and treatment-3 (ambient temperature + 6°C, CO2 750±20ppm). The studied traits are canopy temperature (CT), days to flowering (DtoF), days to physiological maturity (DtoPM), plant height (PH), NDVI, tiller plant^-1^ (TL/P), leaf number (LN), root length (RL), relative leaf water content (RLWC), panicle length (PL), grains panicle^-1^ (G/P), spikelets panicle^-1^ (S/P), spikelet fertility % (SF%), grain yield (GY), thousand-grain weight (TGW) and harvest index (HI).

### 3.5 Selection of genotypes based on yield-associated performance under stress conditions

Genotypic selection was based on the better performance to produce improved yield and yield-associated values with a short phenological span under stress conditions which is comparatively higher than the performance of the check variety - N22 (Figure 6). Based on the less days needed for flowering and higher yield parameters like SB, rice-G/P, SF%, GY and TGW, Kohima special (G2) and Lisem (G6) were found to yielding higher than the N22 (G8). Among the genotypes, Mapok Temeseng (G7) exhibited a longer duration of flowering, with an increase of 33.11% compared to N22. On the other hand, Kaladhan had the shortest duration of flowering (19.94% less compared to N22) (Figure 6A). In case of SF%, Kohima special exhibited the highest increase by 9.48% than the check genotype N22, followed by Lisem which showed a considerable increase in SF (8.09%) (Figure 6D). Also, under stress treatments in both T2 and T3, the performance of these two landraces is much higher comparative to the other landraces. Even in the higher stress condition (T3), trait variables are decreased very less compared to the control and mild stress (T2). Out of the studied eight rice genotypes, Tatza (G1) was found to be a very low yielding landrace under stress conditions. In control, Kohima special and Lisem are yielded 1177.50±17.61 and 1129.67±23.62 g rice grains m^-2^, respectively, which was much higher in range than N22 (1038.50±13.43) and Tatza (814.67±20.48) and upgraded in the mild stress (T2) viz. 1365.50±18.44 for Kohima special, 1294.83±21.73 for Lisem, 1226.67±25.82 for N22 and 859±29.53 for Tatza. But in the severe stress (T3), Kohima special (1108.67±18.61) and Lisem (1062.50±12.74) were performing better than the N22 (985.33±24.72) and Tatza (709.50±34.69) (Figure 6E). Upon analyzing the GY data, a significant difference was observed among the different treatment levels. Kohima special displayed a higher per cent increase in GY (12.32%) than the N22. Lisem also showed notable increases i.e, 7.30% (Figure 6E). When considering all the treatments and compared to the check genotype N22, the landrace Kohima special exhibited the highest increase in TGW by 11.22% (Figure 6F).

**Figure 6.**
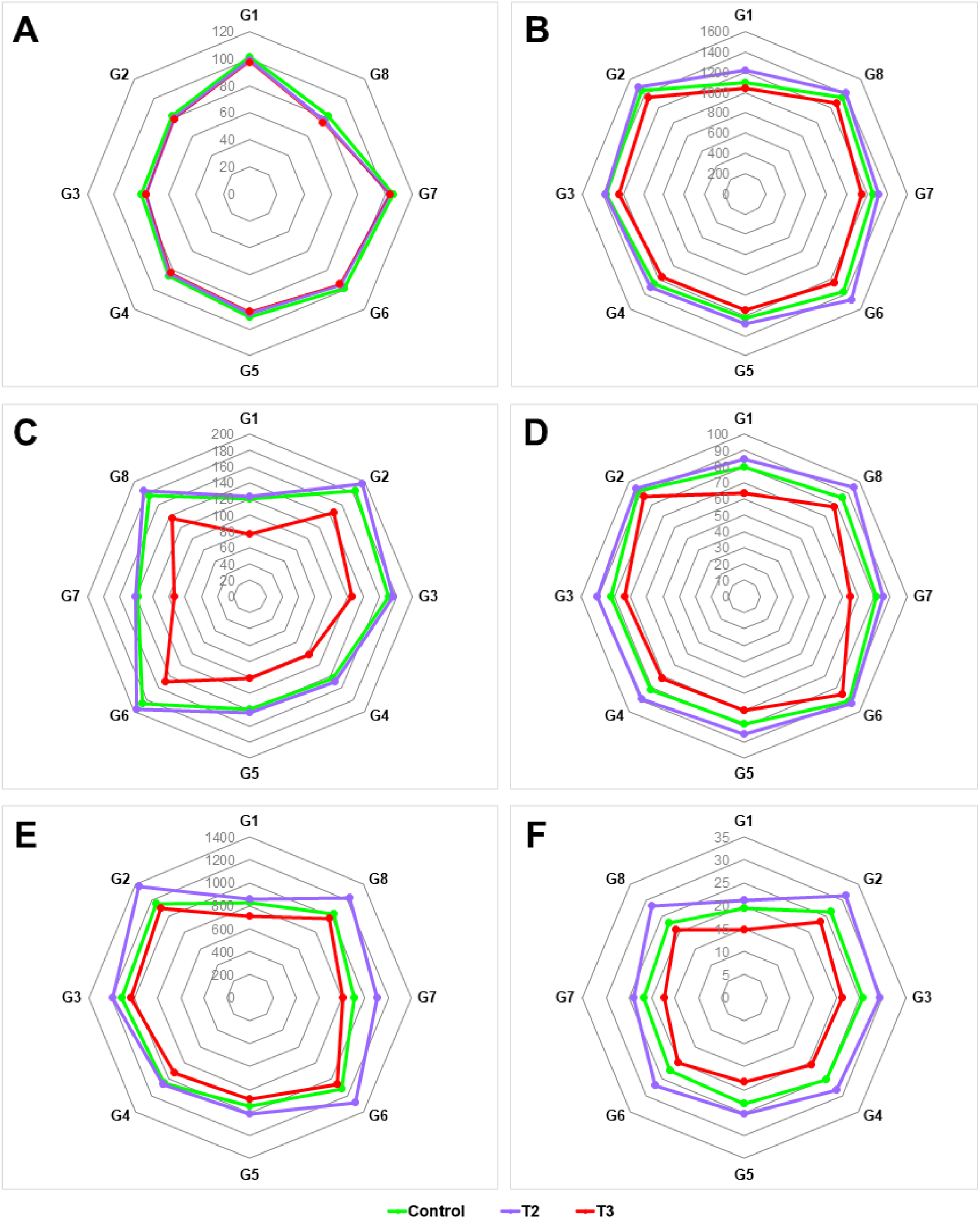
Radar charts showing the mean variability of major yield-associated traits - days to flowering (A), shoot biomass m^-2^ (B), rice-grains panicle^-1^ (C), spikelet fertility% (D), grain yield m^-2^ (E) and thousand-grain weight (F) of studied rice genotypes across two years for the genotypic selection under different high temperature and CO2 level. Genotypes are Tatza (G1), Kohima special (G2), Kaladhan (G3), Laldhan (G4), Tzumma (G5), Lisem (G6), Mapok Temeseng (G7), N22 (G8).

### 3.6 SCoT marker-assisted genetic analysis and genotypic selection

Initially in this study, 30 SCoT markers were screened for genetic analysis among the studied rice genotypes including seven landraces and one check variety. Out of 30 SCoTs, only 25 primers were selected for genetic analysis based on sharp, clear banding patterns and the genotyping information is presented in Table 4. The selected 25 SCoT primers amplified a total of 77 alleles with an average of 3.08 alleles per SCoT locus. One (SCoT28 and SCoT34) to seven (SCoT21) alleles were amplified using the studied marker set that has a 100-1400 bp range of the amplicon size. Out of 77 amplified alleles, 55 were found to be polymorphic with 71.42% of polymorphism. Only eight markers (SCoT5, SCoT9, SCoT15, SCoT19, SCoT20, SCoT21, SCoT31 and SCoT32) were found to be only polymorphic, whereas three (SCoT28, SCoT34, SCoT36) markers were found only monomorphic in nature. The estimates of polymorphic information content (PIC) values ranged from 0.22 to 0.67 with an average of 0.45 per primer. The highest PIC value (0.67) was observed with four alleles in SCoT15. SCoT5, SCoT8, SCoT15, SCoT18, SCoT20, SCoT22, SCoT28, SCoT53, SCoT33 and SCoT35 markers showed PIC value above 0.50. The lowest PIC value (0.22) was obtained with four alleles in the SCoT27 primer (Table 4).

**Table 4.** Molecular genotyping data through 25 SCoT markers-based fingerprinting of rice genotypes including landraces and check variety.

| Sl. No. | SCoT markers | Total amplicons | Amplicon size (bp) | Number of monomorphic bands (%) | Number of polymorphic bands (%) | PIC<sup>1</sup> |
| --- | --- | --- | --- | --- | --- | --- |
| 1 | SCoT 2 | 2 | 180-300 | 1 (50%) | 1 (50%) | 0.44 |
| 2 | SCoT 5 | 3 | 100-400 | 0 | 3 (100%) | 0.52 |
| 3 | SCoT 6 | 6 | 150-750 | 1 (16.66%) | 5 (83.33%) | 0.38 |
| 4 | SCoT 8 | 4 | 300-800 | 2 (50%) | 2 (50%) | 0.58 |
| 5 | SCoT 9 | 2 | 140-190 | 0 | 2 (100%) | 0.30 |
| 6 | SCoT15 | 4 | 190-1050 | 0 | 4 (100%) | 0.67 |
| 7 | SCoT16 | 3 | 160-480 | 1 (33.33%) | 2 (66.66%) | 0.50 |
| 8 | SCoT17 | 4 | 110-820 | 1 (25%) | 3 (75%) | 0.43 |
| 9 | SCoT18 | 3 | 900-1400 | 1 (33.33%) | 2 (66.66%) | 0.60 |
| 10 | SCoT19 | 2 | 120-260 | 0 | 2 (100%) | 0.49 |
| 11 | SCoT20 | 2 | 170-350 | 0 | 2 (100%) | 0.55 |
| 12 | SCoT21 | 7 | 220-900 | 0 | 7 (100%) | 0.37 |
| 13 | SCoT22 | 4 | 400-1000 | 1 (25%) | 3 (75%) | 0.58 |
| 14 | SCoT24 | 3 | 400-1200 | 2 (66.66%) | 1 (33.33%) | 0.34 |
| 15 | SCoT26 | 3 | 400-1250 | 2 (66.66%) | 1 (33.33%) | 0.38 |
| 16 | SCoT27 | 4 | 200-550 | 1 (25%) | 3 (75%) | 0.22 |
| 17 | SCoT28 | 1 | 1000 | 1 (100%) | 0 | 0.52 |
| 18 | SCoT29 | 4 | 300-1200 | 2 (50%) | 2 (50%) | 0.47 |
| 19 | SCoT30 | 4 | 200-800 | 1 (25%) | 3 (75%) | 0.53 |
| 20 | SCoT31 | 2 | 300-500 | 0 | 2 (100%) | 0.29 |
| 21 | SCoT32 | 2 | 120-280 | 0 | 2 (100%) | 0.38 |
| 22 | SCoT33 | 3 | 150-500 | 1 (33.33%) | 2 (66.66%) | 0.58 |
| 23 | SCoT34 | 1 | 800 | 1 (100%) | 0 | 0.30 |
| 24 | SCoT35 | 2 | 180-500 | 1 (50%) | 1 (50%) | 0.57 |
| 25 | SCoT36 | 2 | 300-600 | 2 (100%) | 0 | 0.33 |
| **Total** |  | **77** |  | **22** | **55** |  |
| **Average** |  | **3.08** |  | **0.88** | **2.2** | **0.45** |

<sup>1</sup>Polymorphism information content (PIC)

The dendrogram, based on SCoT-based genotyping data, classified the studied rice genotypes into two broad clusters – I and II (Figure 7). Cluster I comprise six landraces – Tatza (G1), Kohima special (G2), Kaladhan (G3), Laldhan (G4), Lisem (G6) and Mapok Temeseng (G7), whereas cluster II comprises two genotypes – one landrace i.e. Tzumma (G5) and one modern check variety i.e. N22 (G8). In cluster I, Jaccard’s similarity coefficient shows the highest similarity (75%) between Kohima special and Lisem and they are connected closely to each other. Then they sub-clustered with Kaladhan, which further sub-clustered with Laldhan and Mapok Temeseng. These all five genotypes then distantly clustered with Tatza.

**Figure 7.**
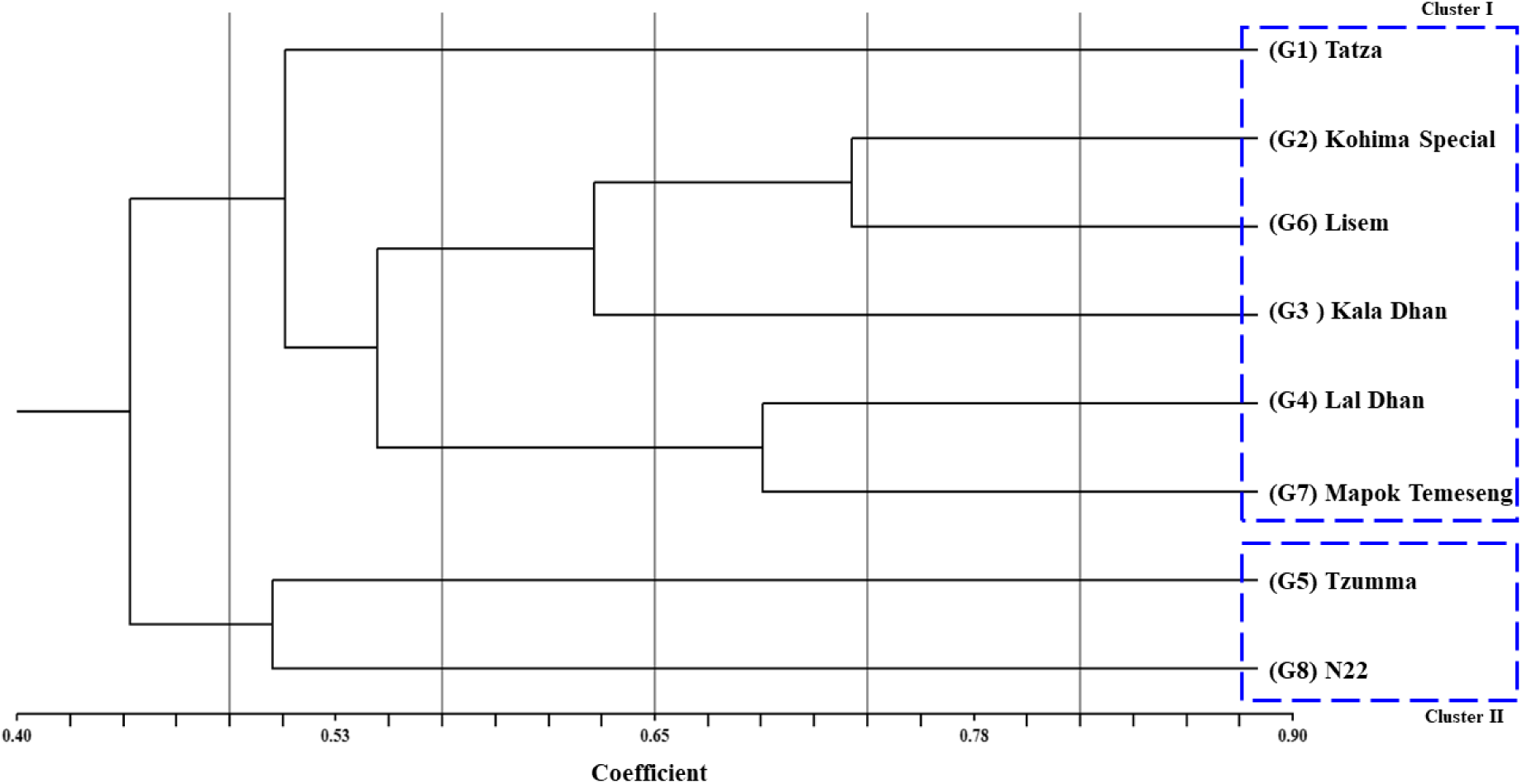
Dendrogram showing the relationship between studied rice genotypes using SCoT marker-based genotyping data

## 4. Discussions

The present study sheds light on the potentiality of rice landraces of the North-East Indian region to perform better than the commonly used check variety under high temperatures and elevated CO_2_ levels in the context of the present day’s ongoing climate change. The observed variability in terms of phenological, morphological, physiological, and yield-associated traits in the studied genotypes under mild and severe treatments of high-temperature and elevated CO_2_ in two successive cropping seasonal years (2019 and 2020) were further confirmed with SCoT marker-based genetic profiling.

Rice landraces, especially from the North-East Indian location are important pre-breeding materials for future rice breeding programs. Numerous studies have demonstrated that indigenous rice varieties that farmers have year-long cultivated, maintained, and preserved in their local agro-ecological niche have a high level of genetic diversity, making them viable genetic resources for enhancing yield, agronomic performance and tolerance to abiotic stresses (Choudhury et al., 2013; Pusadee et al., 2009). Evolutionary, landraces play a bridging role in domestication from wild species to modern high-yielding rice varieties. In this investigation, the initial screening led to identify seven comparatively heat-tolerant rice landraces out of a group of 75 population which is very easy and well-performing during the big trial in field conditions. Such screening was based on the visible changes caused by the high temperature – like leaf and leaf-tip rolling, curling and rice plant death (Table S1; (Hasanuzzaman et al., 2013). For further analysis, a national check rice variety Nagina22 (or N22) developed by the National Rice Research Institute (NRRI, Cuttack, India) was added. Many other studies used this heat-tolerant variety for a comparative study of heat stress response by other rice genotypes (Behera et al., 2023; Kilasi et al., 2018).

Evaluation of the genotypes under stress treatments reveals the profound effects of the high temperatures and elevated CO_2_ levels on rice phenology, morpho-physiological and yield-associated parameters (Table 2). Year-wise variability of almost every trait was very low because the traits were mostly genetically stable through the year-wise environmental fluctuations. So, further deviation must come from the treatments and genotypes. A noticeable pattern was seen wherein the mild stress treatment (T2) of high temperature (ambient + 4°C) and 550 ppm CO_2_ resulted in higher trait values compared to the control. However, this trend significantly diminished under much higher temperatures (ambient + 6°C) with higher CO_2_ (750 ppm) levels (T3). The stress-induced reduction of growth parameters was greatly ameliorated by elevated CO_2_. Elevated CO_2_ with high temperature shows a significant difference in plant height among the genotypes in both years. The reduction in DtoF and DtoPM in mild stress is of great importance in shortening the life cycle, which may result in higher yields under stressful conditions for the tolerant genotypes (Bheemanahalli et al., 2017). Reduction in DtoF and increment of yield-associated traits in rice in response to high temperature and elevated CO_2_ conditions, respectively, has been reported in other studies (Raj et al., 2016). Early flowering and maturity are important to minimize the cost-effective agricultural input during the cropping season and may help to avoid the onset of climatic instabilities (Bhaduri et al., 2023). The treatment effect was visualized in most of the yield-associated traits as they are multigenic and very hard to regulate. For reproductive success, pollen (male reproductive particle) viability, P/A and AnL are crucial for higher yield (Roychowdhury et al., 2012). Such pollen traits are reported a significant difference in our previous study on hot chilli grown under high temperatures and elevated CO_2_ (Das and Das, 2021). It has been noted the effect of high temperature on pollen viability in rice grown under polyhouse with elevated temperature from 29°C to 40°C and maintained until 15 h, with a relative humidity of 75%. Such experimental set-up exhibited 20 - 40% SF after exposure to high temperature. Moreover, the pollen fertility ranged from 39 to 90% (Jagadish et al., 2007). In order to validate the contribution of AnL and P/A to SF% at maturity, both found a significantly positive correlation in different stress levels (Figure 2A-B), but the strength was much higher between P/A and SF% as because pollens in anther has a direct influence on rice reproductive success, which in turn may eventually provide higher grain set panicle^-1^ and higher GY (Figure 2C). Following our results, (Sakai et al., 2019) also found that elevated CO_2_ increases the number of SP/P and grain set. An earlier study reported a higher TL number, LA and SB (above ground) of hybrid rice cultivars under elevated CO_2_ conditions. The higher biomass can be also related to a higher N_2_ supply and a minimum (1-2°C) temperature elevation (Hu et al., 2022). The root is also an important stress-responsive plant part which uptake nutrients from the soil and channelizes to shoot and leaf to influence plant growth and development. The photo-assimilates are stored in shoot and root parts and improve their biomass. Under abiotic stress conditions, there is a considerable effect on rice roots especially root length, number, branching and volume (Roychowdhury et al., 2013; Yang et al., 2007). There was a notable difference in root length among the genotypes in both years. (Jin et al., 2019) reported that improved root traits under elevated CO_2_ were due to increased photosynthetic carbon allocation towards roots which stimulates root growth and thereby enhances water and nutrients uptake efficiency. As a consequence, RB and root : shoot ratio in rice increased in response to higher temperatures (Kim and You, 2010). In many cereal crops including rice, it has been tested and proved that higher biomass can increase GY and grain set through channeling higher nutrients and photo-assimilates to the growing panicles during grain set (Roychowdhury et al., 2023b). In the present study, SB significantly and positively correlates with GY, SP/P, G/P, HI, and TGW in stress treatments (Figure 2D-I) except PL (non-significant associations) which is showing very less R^2^ value i.e. 0.76 (Table 3). Generally, it may seem long panicle can hold higher grains for yielding high, but the finding indicates higher advantages of panicle density or grain set panicle^-1^ on the yield rather than PL. The probable cause may be in the mild stress conditions, CO_2_ level helps to accelerate photosynthesis rate due to higher LN or greater LA to assimilate higher carbon to improve biomass, which is eventually boosting GY (Kim and You, 2010). We have found evidence of higher LA corresponds to P/A and GY in this study (Figure 4B).

Under high temperatures and elevated CO_2_ stress conditions, the physiological traits of rice play a very crucial role in characterizing genotypes and stress effects. In our study, CT, chlorophyll content (SPAD), NDVI, RL, RLWC, and LA are found significantly correlated with GY (Figure 3A-F). Stress-tolerant rice plants have a characteristic feature to adjust their body temperature lower by exposure to stress environment and internally can function in stress-resilient ways to produce higher grain production and/or minimize the trade-off of yield penalties because of harsh stress (VanWallendael et al., 2019). In this case, CT is negatively associated with the GY increase, whereas the other physiological characteristics are positively associated with GY. NDVI also indicates the vegetation and biomass of the crop and thus shows a significant correlation (r=0.94) with G/P (Figure 4A). As was discussed earlier that years (Y) has the very least effect on the traits as most of the traits show significant differences in the case of genotypes, treatments, and years individually, but in some instances where G and T are significant effects, years is not producing significant variable differences for DtoF, DtoPM, AnL, RB, TL m^-2^ and LN (Table 3). Multivariate analysis exhibits a clear separation of treatments, mainly the severe stress condition is distinctly separated in the PCA plane due to its harsh negative effect on the studied traits, whereas control and mild stress are almost mingled in the place because of their similar or close effect (Figure 5A1-B1). In most of the traits, mild stress has a beneficial effect over the control as we stated before. Phenology is grouped in the close vicinity of CT. However, the plant height vector is closely situated with other physiological and yield-associated traits. In general, for the modern varieties, PH is inversely proportional to the GY as a resolution after the Green Revolution. But as we have used most of the landraces, it represents higher PH-mediated biomass increment and higher yielding. When considering selecting the best-yielding rice landraces that have similar or higher performance than the national check variety under stress conditions, Kohima special (G2) and Lisem (G6) have been identified as potent landraces which have comparatively shorter DtoF and higher SB, grain set panicle^-1^, SF%, GY and TGW (Figure 6).

In our investigation, SCoT markers were used to assess the molecular effect of high temperature and elevated CO_2_ on genotypes’ genetic diversity. In other abiotic stress responsiveness of rice, some other molecular markers like SSR, ISSR, SNP were used in the rice landrace population (Ganie et al., 2016, 2014; Karmakar et al., 2012; Meena et al., 2023; Roychowdhury et al., 2013). SCoT marker-based analysis is better than other frequently used molecular markers since primer creation does not need genomic sequence information (Omar et al., 2023)(Omar et al. 2023). SCoT markers also have larger polymorphism rates than SSRs, making them better at distinguishing closely related genotypes. SCoT analysis eliminates the resource-intensive SNP identification and genotyping, making it cheaper than SNP markers. SCoT markers also target non-coding areas around start codons, which may identify gene expression and functional variation regulators (Patidar et al., 2022; Que et al., 2014; Zhang et al., 2015). In our study, eight SCoT markers (SCoT5, SCoT9, SCoT15, SCoT19, SCoT20, SCoT21, SCoT31 and SCoT32) have indented highly polymorphic and can be used further for rice genotyping for high temperature and CO_2_ induced stress. The polymorphism information content (PIC) of the studied SCoT markers is moderate and signifies each marker’s genotypic discrimination ability. The best performing two rice landraces (Kohima special and Lisem) under high temperature and elevated CO_2_ level were obtained from multivariate analysis, which is supported by the SCoT marker-based selection data as showing the similarity in marker-based clustering (Figure 7).

In such a way, rice landraces exhibiting resilience and superior performance under high temperatures and elevated atmospheric CO_2_ stress conditions should be prioritized under ongoing climate change scenarios in agricultural sectors. However, stability over multiple environments and seasons is critical, so multi-location trials should be conducted to validate their adaptability. Additionally, assessing the genetic diversity present in the selected rice landraces will guide breeders in avoiding narrow genetic bases and potential vulnerabilities. New climate-resistant rice varieties may be developed using the genetic diversity of Kohima special and Lisem. These landraces can provide sustainable rice production in the face of climate change by incorporating traits like efficient photosynthesis, heat tolerance, and water-use efficiency. This will improve food security for local communities and the wider population.

## 5. Conclusion and future breeding scope

The present investigation employed the evaluation of rice landraces for better performance under high temperatures and elevated CO_2_ for breeding traits and further aided with SCoT marker-assisted selection. By comprehensively assessing morphological, phenological, physiological and yield-associated traits, this finding led to identify two potent rice landraces – Kohima special and Lisem having higher yielding ability under stress conditions, much superior than the national check variety N22. SCoT marker-based genotyping also revealed moderate genetic diversity among the genotypes and found few potent markers (SCoT5, SCoT9, SCoT15, SCoT19, SCoT20, SCoT21, SCoT31 and SCoT32) with better discrimination ability to precisely select the superior genotypes that would help the rice breeders in selection of suitable parents for breeding purpose and genetic mapping studies. By using these two landraces as donor parents, the genetic background of N22 or other modern cultivars can be improved to sustain in the changing climatic scenarios. The findings could influence rice breeding programs, guiding the selection of rice varieties that are better equipped to thrive in the changing environmental conditions. This multidimensional perspective can shape the future of rice breeding and contribute to global efforts aimed at achieving food security in an era of environmental uncertainty.

## Abbreviations

AnL: anther length
CT: canopy temperature
DtoF: Days to flowering
DtoPM: Days to physiological maturity
G/P: grains panicle^-1^
GY: grain yield
HI: harvest index
LA: leaf area
LN: leaf number
NDVI: normalized difference vegetation index
P/A: pollens per anther^-1^
PCA: principal component analysis
PH: plant height
PL: panicle length
RL: root length
RLWC: relative leaf water content
RV: root volume
SB: shoot biomass
SF%: spikelet fertility %
SP/P: spikelets panicle^-1^
SPAD: soil-plant analysis development
TGW: thousand-grain weight
TL/P: tillers per plant

## Acknowledgements

The authors are grateful to the Assam Agricultural University (AAU, Jorhat, India) for providing the resources for conducting the experiments and support this present research.

## CRediT authorship contribution statement

**RD** conceived the proposal, supervised the work and funding acquisition; **MM** collected resources; **MM**, **RR**, **RS**, **LT** and **DJ** performed the investigation, experiments, sampling and data collections; **RR**, **RSS** and **DJ** analyzed the data and visualization through software; **MM**, **RD** and **RR** worked on methodologies; **MM**, **RS** and **RD** curated the data; **MM**, **RR** and **RS** wrote the original manuscript; **BG**, **PK**, **DJ** and **RD** reviewed and edited the manuscript. All authors have read and agreed to the published version of the manuscript.

---

**Table S1.**
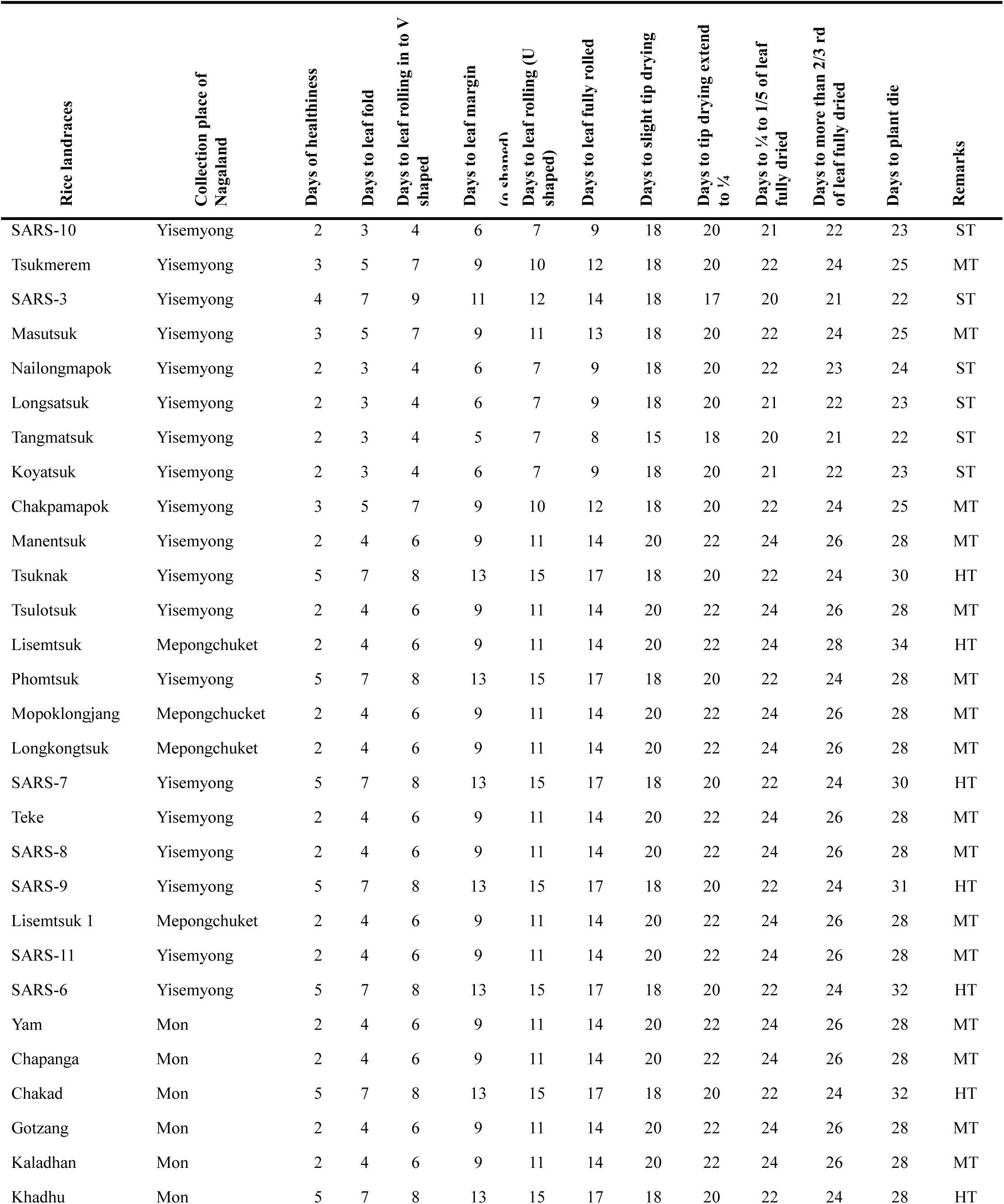
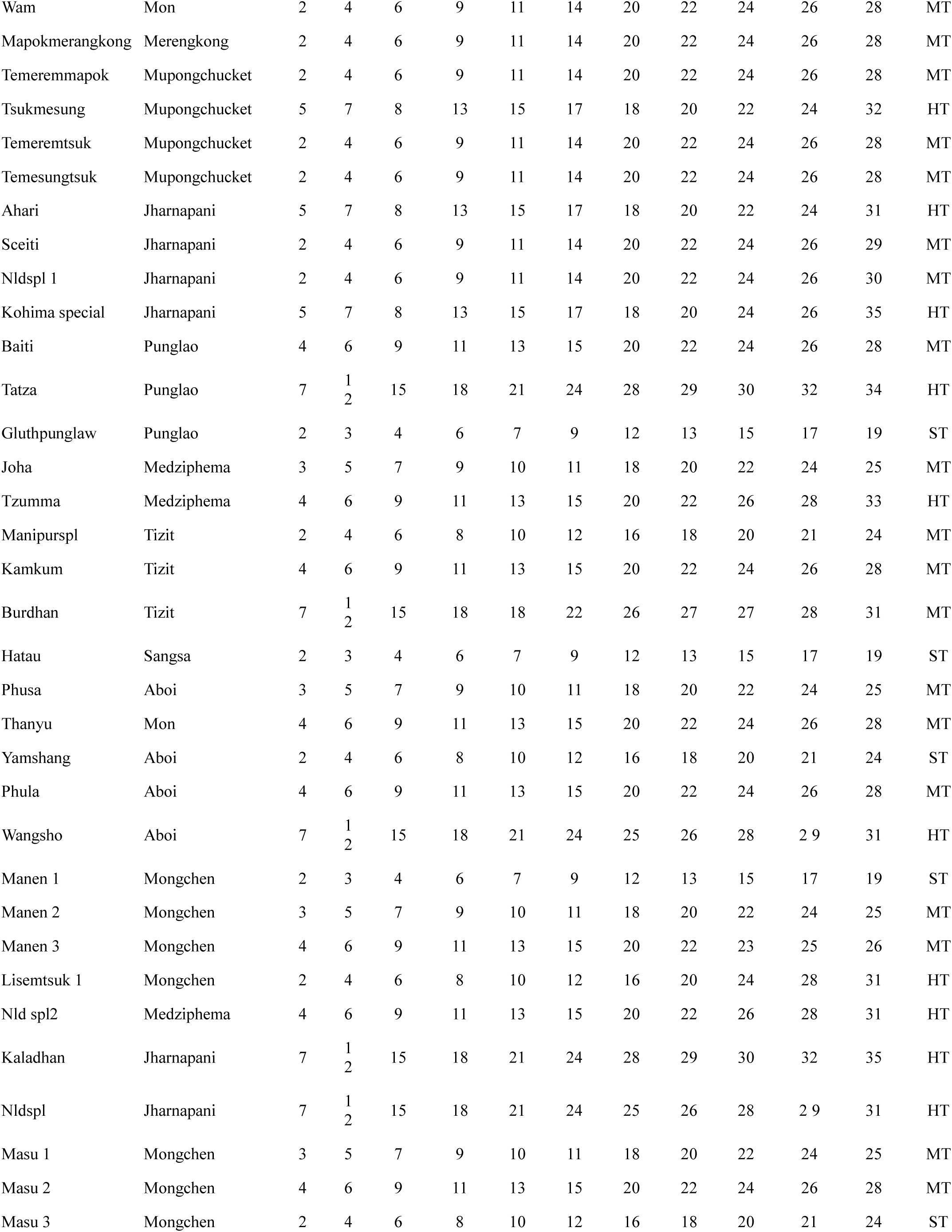
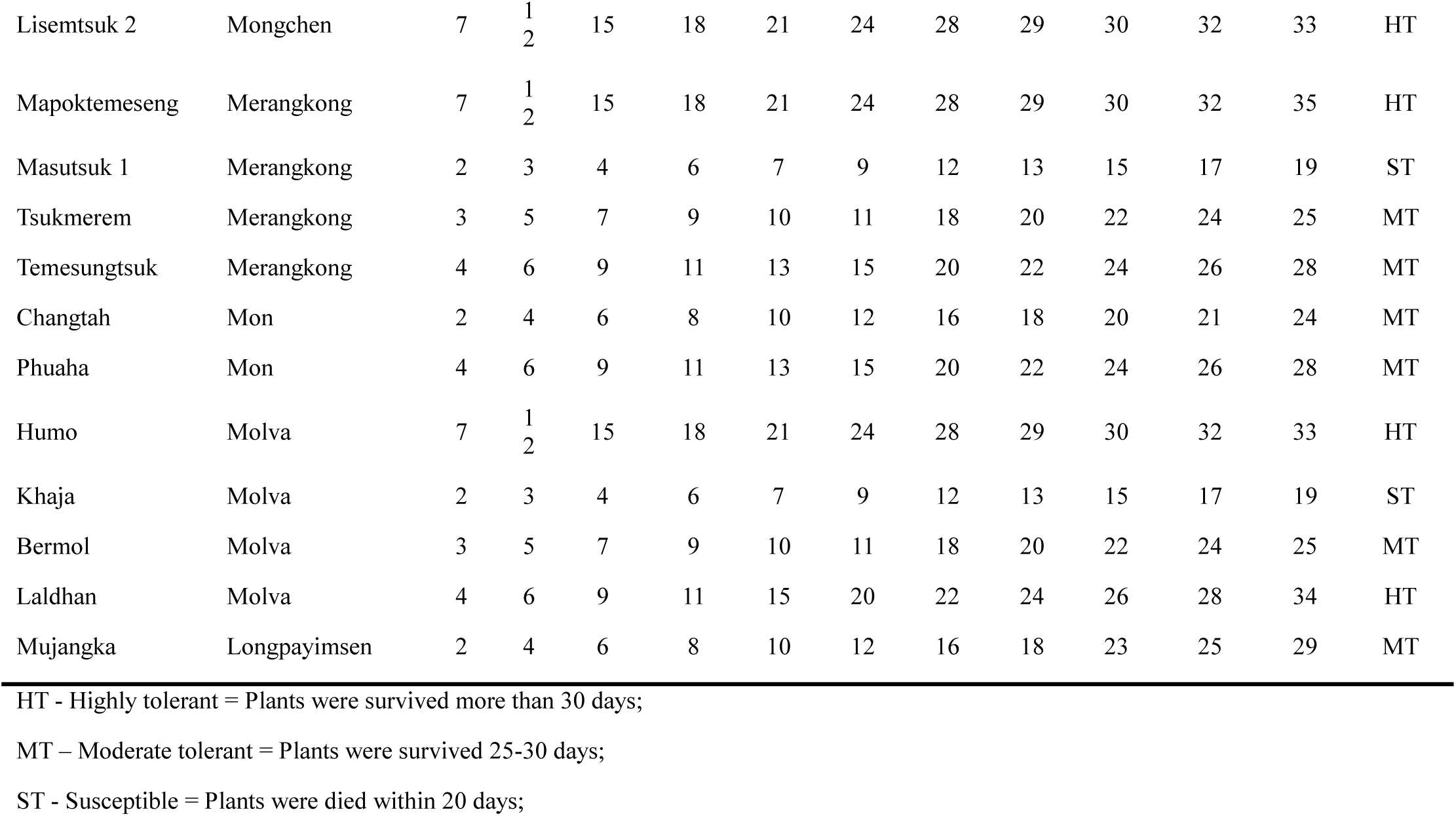
Screening of rice landraces from North East India (Nagaland) under high temperature stress condition.

**Table S2.**
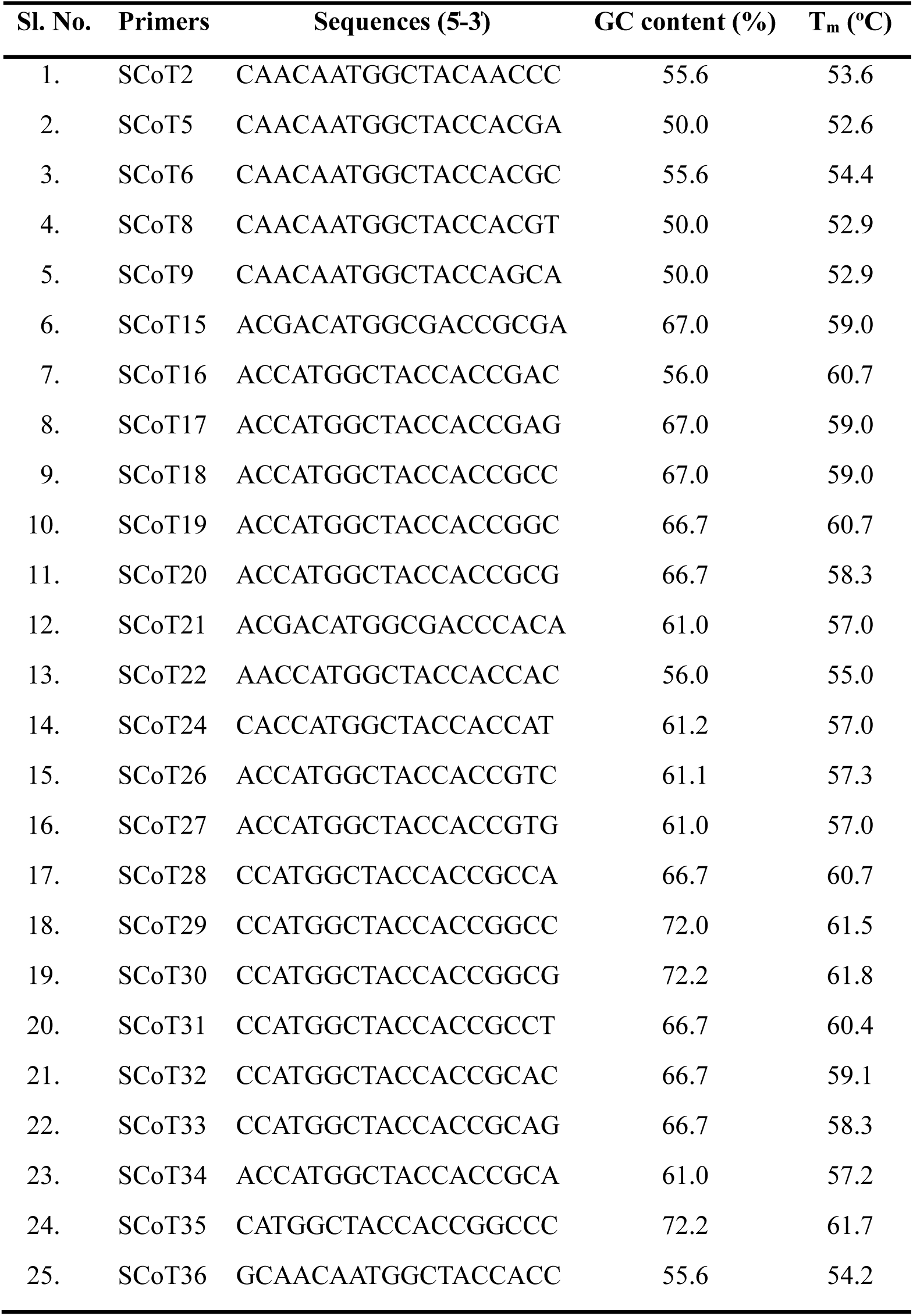
Details of used SCoT primers and their sequences, GC content and melting temperature (Tm)

